# LINC01133 knockout increases malignancy by migration mechanisms in Hs578T Triple-Negative Breast Cancer Cells

**DOI:** 10.64898/2026.07.03.736417

**Authors:** Henrique César Jesus-Ferreira, Leandro Teodoro, Ana Claudia Oliveira Carreira, Mari Cleide Sogayar

## Abstract

Long non-coding RNAs (lncRNAs) have attracted increasing interest because of their roles as modulators of tumor progression, acting either as oncogenic drivers or tumor suppressors, depending on the cellular context. LINC01133 has been implicated in regulation of multiple tumor-related mechanisms; however, its role in breast cancer, particularly in the triple-negative subtype, remains poorly characterized. In this study, we investigated the impact of LINC01133 depletion on malignant phenotypes and on the expression of migration- and invasion-associated genes using the Hs578T triple-negative breast cancer (TNBC) cell line, through comparative analyses of parental, control, and LINC01133-knockout cell lines, namely Hs578T_wt, Hs578T_ctr, and Hs578T_ko. Functional characterization included morphological analysis, growth assays, anchorage-independent colony formation, migration, invasion, and quantitative biomolecular experiments. Depletion of LINC01133 led to reduction of cell diameter, a significant increase in colony-forming capacity, and marked enhancement of migratory and invasive potential. At the molecular level, LINC01133 loss induced the expression of genes associated with extracellular matrix remodeling and cellular plasticity, including fibronectin, vimentin, integrins, FOXC1, and TWIST1, concomitant with reduced expression of ZEB1, TWIST2, and N-cadherin. Collectively, these data indicate that LINC01133 acts as a potential fine regulator of *in vitro* migration and invasion processes in TNBC, with its expression favoring a more asymptomatic mode of tumor progression, whereas its loss markedly enhances tumor malignancy.

## 1. Introduction

TNBC is recognized as the most aggressive molecular subtype of breast cancer, being associated with high mortality rates, particularly in cases of recurrence, since only two out of ten women survive [1]. High lethality is associated with a set of intrinsic and extrinsic tumor factors, among which are the absence or low expression of Estrogen Receptor (ER), Progesterone Receptor (PR), and Human Epidermal Growth Factor Receptor-type 2 (HER2), which are biomarkers traditionally explored as therapeutic targets in other breast cancer subtypes [2]. The lack of these targets significantly limits treatment options, rendering systemic chemotherapy the main clinical approach available, although it often has restricted efficacy and is associated with early and aggressive relapses.

TNBC is characterized by pronounced molecular heterogeneity, reflected in the existence of at least six subclasses with distinct tumor progression profiles. The Basal-like 1 and Basal-like 2 (BL1 and BL2, respectively) subclasses are characterized by strong proliferative activity; the Mesenchymal (M) and Mesenchymal Stem-like (MSL) subclasses are associated with migratory and mesenchymal phenotypes, the Immunomodulatory (IM) subclass presents an inflammatory profile with enrichment in immune-related signals, and the Luminal Androgen Receptor (LAR) subclass is distinguished by an androgenic transcriptional signature. This phenotypic and molecular diversity contributes not only to clinical variability but also to therapeutic resistance and tumor adaptation to microenvironmental pressures [3,4].

Another crucial aspect lies in the intratumoral environment of TNBC, since fine regulation of intra- and intercellular signaling pathways sustains tumor plasticity. The tumor microenvironment (TME) is highly dynamic and complex, integrating cellular components (cancer-associated fibroblasts, immune cells, and endothelial cells) and non-cellular components (extracellular matrix, ECM, and soluble factors) into interactions that shape tumor behavior [5]. At this interface, gene-switch mechanisms play a fundamental role by promoting ECM remodeling and regulating its stiffness and structural composition through the differential expression of structural genes and crosslinking proteins. These processes contribute not only to the generation and maintenance of Cancer Stem Cells (CSCs), but, also, to the activation of effective mechanisms of immune evasion, which supports tumor progression and delays immune recognition [6].

This molecular and microenvironmental complexity makes elucidating the fundamental mechanisms involved in TNBC progression challenging. In this context, several non-coding RNAs (ncRNAs) have emerged as key elements, since they modulate gene expression through post-transcriptional mechanisms or by direct interaction with regulatory proteins [7]. Among ncRNAs, the long non-coding RNAs (lncRNAs) play a prominent role, with multiple lines of evidence pointing to their active involvement in the regulation of critical pathways of tumorigenesis, tumor progression, cell migration, invasion, and microenvironment remodeling [8].

A particularly relevant example is LINC01133, a member of the lncRNA subclass. This transcript, originating from a homonymous gene, has multiple isoforms and has been predominantly associated with tumor suppressor functions in different contexts. Structurally, LINC01133 gene contains three canonical exons and four internal loops, which, according to in silico analyses, may act as miRNA sponges and protein interaction scaffolds, suggesting a central role in regulation of molecular networks [9].

In breast cancer, LINC01133 has been described as a tumor suppressor. Recent studies have demonstrated its activity in human TNBC cell lines, such as MDA-MB-231 and Hs578T, relating LINC01133 with its role in mesenchymal cells present in the TME, its capacity to act as a sponge for miR-199a, and potential associations with regulatory proteins, such as KLF4 and EZH2, which remain incompletely characterized [10,11]. Despite these advances, the molecular mechanisms underlying its function in TNBC remain largely unknown, requiring further investigation.

Considering the heterogeneity of expression and multiple potential interactions of LINC01133 in intratumoral and microenvironmental models of breast cancer, in the present study, we explore the biological functions of LINC01133 in the Hs578T cell line model. To this end, LINC01133 gene knockout was established, accompanied by functional phenotypic analyses. The results indicated an increase in malignancy in the knockout modified cell line, with an emphasis on phenotypes related to tumor migration and invasion. Therefore, we propose that LINC01133 may act as a tumor suppressor lncRNA, modulating pathways related to epithelial-mesenchymal transition (EMT), cell adhesion, and tumor invasion. Consequently, its functional loss may trigger a more aggressive phenotype, favoring enhanced migratory and invasive phenotype. This supports the importance of investigating this lncRNA as a potential molecule of interest in TNBC dynamics.

## 2. Materials and Methods

### 2.1. Cell lines descriptions

The following human cell lines were used in this study: MCF10A (RRID:CVCL_0598), a non-tumorigenic mammary epithelial cell line; MCF12A (RRID: CVCL_3744), a non-tumorigenic mammary epithelial cell line; MCF7 (RRID:CVCL_0031), a luminal A breast adenocarcinoma cell line; ZR-75-1 (RRID:CVCL_0588), a luminal B breast ductal carcinoma cell l ine; SK-BR-3 (RRID:CVCL_0033), a HER2-positive breast adenocarcinoma cell line; MDA-MB-231 (RRID:CVCL_0062), a basal-like, triple-negative breast adenocarcinoma cell line; and Hs578T (RRID:CVCL_0332), a basal-like, triple-negative breast carcinoma cell line derived from mammary gland tissue. All cell lines were obtained from the American Type Culture Collection (ATCC, Manassas, VA, USA).

Cell line authentication was performed following the test recommendations provided by American Type Culture Collection (ATCC) (Cell Line Authentication Testing Recommendations). Authentication was conducted by short tandem repeat (STR) profiling, and the resulting profiles were compared with reference databases to confirm cell identity and exclude cross-contamination. Only cell cultures showing ≥ 80% concordance with reference STR profiles were used in this study. Experiments were performed using cells maintained for no more than 15 passages after thawing. All cell lines were routinely tested for mycoplasma contamination using a nested PCR assay standardized by our group. Only mycoplasma-free cultures were used in all experimental procedures.

The Hs578T cell line was used to generate two genetically modified cell lines through lentiviral transfection followed by transduction of the Hs578T_wt. The cell line carrying an inserted closed, empty plasmid vector, used as a control for the genetic modification process, was designated as the control line (Hs578T_ctr), while the cell line in which LINC01133 gene was knocked out has been designated as the LINC01133 knockout cell line (Hs578T_ko). The experimental procedures for generation and validation of these cell lines were described in detail by Teodoro Júnior et al. (2026) [12] and Jesus-Ferreira (2021) [13].

### 2.2. Specifications of Cell cultures

All parental cell lines used were obtained from the ATCC (Virginia, USA). MCF10A and MCF12A cell lines were cultured in DMEM/F12 1:1 [+] L-Glutamine, [+] Bicarbonate, [+] Phenol-Red (Sigma-Aldrich, St. Louis, MO, USA) supplemented with 10% Fetal Bovine Serum (FBS) (Athena, São Paulo, Brazil), EGF (20 ng/mL, Gibco, Grand Island, NY, USA), Hydrocortisone (0.5 mg/mL, St. Louis, MO, USA), and Insulin (10 µg/mL, Sigma-Aldrich, St. Louis, MO, USA); MCF7 cell line was cultured in EMEM (Sigma-Aldrich, St. Louis, MO, USA) supplemented with 10% FBS and Insulin (Sigma-Aldrich, 4 mg/mL); the ZR-75-1 strain was cultured with RPMI-1640 culture medium (Sigma-Aldrich, St. Louis, MO, USA) supplemented with 10% FBS (Athena, São Paulo, Brazil); the SK-BR-3 strain was cultured with McCoy’s 5A culture medium (Sigma-Aldrich, St. Louis, MO, USA) supplemented with 10% FBS; the MDA-MB-231 and Hs578T cell lines were cultured with DMEM High Glucose, [+] Phenol-Red (Sigma-Aldrich, St. Louis, MO, USA) supplemented with 10% FBS (Athena, São Paulo, Brazil). All media were supplemented with the antibiotics Ampicillin (25 mg/L) and Streptomycin (100 mg/L) (Sigma-Aldrich, St. Louis, MO, USA).

### 2.3. lncRNAs qRT-PCR panel expression of cell lines

All parental cell lines were cultured in medium-sized flasks (T75, Corning, NY, USA) until reaching 90% confluence. Total RNA was then extracted from 1×10⁶ cells using the RNEasy Mini Kit (QIAGEN, Hilden, Germany). cDNA synthesis was performed with 2 µg of RNA from each experimental condition using the FireScript RT cDNA Synthesis Kit (Solis Bio-Dyne, Tartu, Estonia). The yield and purity of the synthesized cDNA was assessed by the 280/260 nm ratio, with A260/A280 ratio above 2.0, using a Nanodrop 1000 (Thermo-Scientific, Waltham, MA, USA), according to the manufacturer’s instructions.

For qRT-PCR, the Fast SYBR™ Green Master Mix kit (Cat #4385612, ThermoFischer, Waltham, MA, USA) was used with a 40-cycle amplification protocol following the basic recommendations of the equipment (ViiA7 Real-Time PCR System, ThermoFischer, Waltham, MA, USA). Each reaction contained 2.5 µL of total cDNA, 2.5 µL of primers (forward + reverse, 600 nM) for the lncRNAs target genes (or 2.5 µL of nuclease-free H₂O in negative-control wells), and 5.0 µL of Fast SYBR™ Green Master Mix (optimized volumes). Primers were obtained from Exxtend – *Soluções em Oligos* (Paulínia, São Paulo, Brazil), lyophilized and desalted, as listed in the table below (Table 1):

**Table 1:**
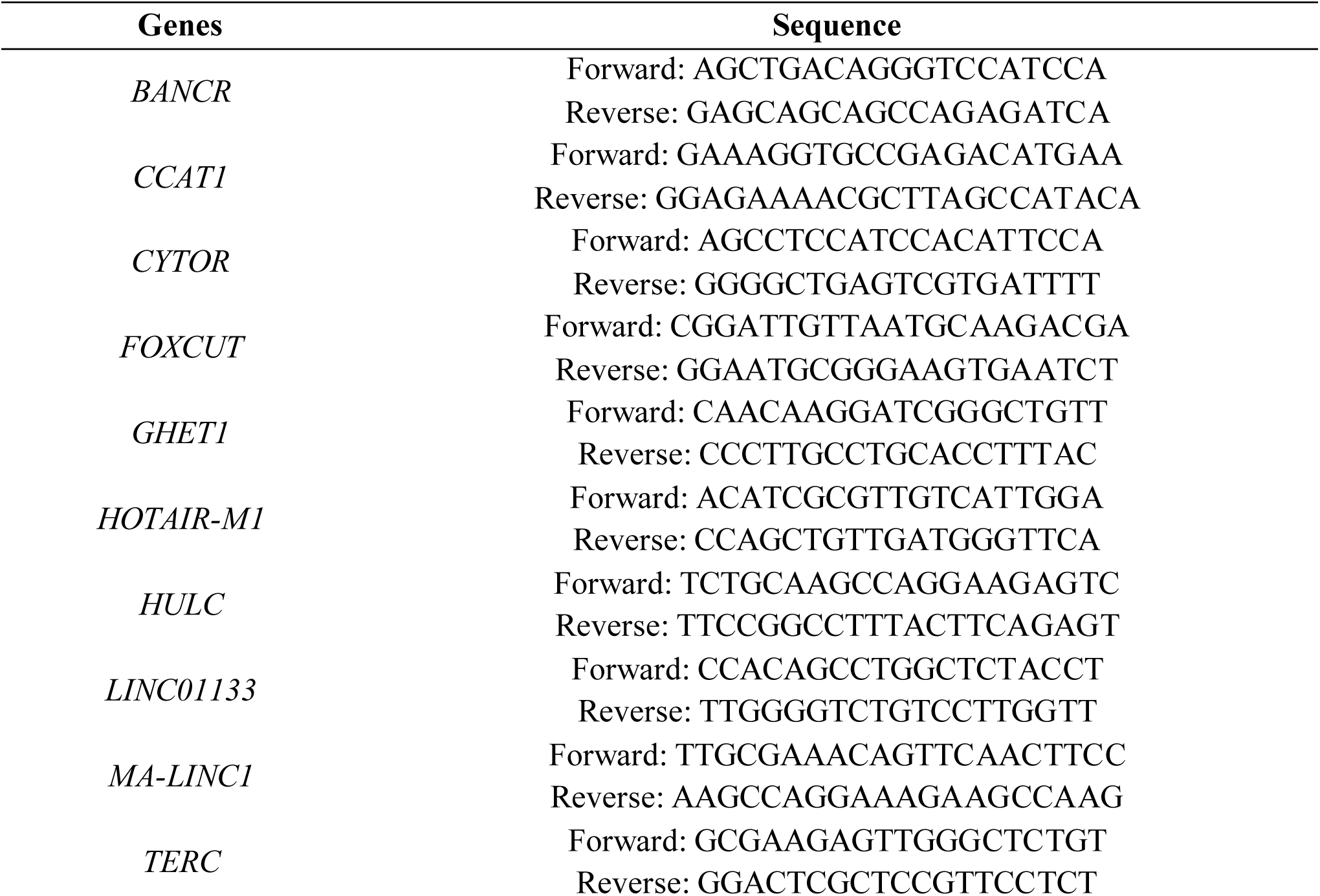
Primers sequences of lncRNAs genes analyzed.

The internal control (reference) genes used were HPRT1 (600 nM, Forward: TGAGGATTTGGAAAAGGGTCT; Reverse: GAGCACACAGAGGGCTACAA) and HMBS (600 nM, Forward: TGGACCTGGTTGTTCACTCCTT; Reverse: CAACAGCATCATGA GGGTTTT).

All qRT-PCR assays were performed in technical and biological triplicate. The mean Ct values of the internal control genes were used to calculate the log₂FC values for each analyzed gene using the 2^−ΔΔCt^ method (reference cell line: MCF10A). Final results were analyzed statistically using one-way ANOVA with a significance threshold of p ≤ 0.05 (GraphPad Prism, v. 10.6, Boston, MA, USA).

### 2.4. Growth curves and phenotypic assays

For the growth curve assays and subsequent experiments, the Hs578T_wt, Hs578T_ctr, and Hs578T_ko cell lines were used. For all assays, cells were cultured until reaching 80% confluence to maintain higher mitotic potential [14]. For the growth curve experiments, cells in culture were subcultured into 35 mm culture dishes (Corning, NY, USA), in technical triplicates and biological duplicates, under three growth conditions: 10%, 5%, and 1% FBS, at a density of 1×10⁴ cells per dish. Cell growth was monitored over 11 days, with cell collections (harvesting) on days 1, 3, 5, 7, 9, and 11, and medium changes every three days. Cell collection followed a standard trypsinization protocol (Trypsin, Sigma-Aldrich, St. Louis, MO, USA) and fixation with 4% v/v PFA in PBSA (PFA, Sigma-Aldrich; PBSA, Sigma-Aldrich, St. Louis, MO, USA). Cells were quantified by flow cytometry using an Accuri C6 Plus Flow Cytometer (BD Biosciences, San Jose, CA, USA), based on FSC-H and SSC-H parameters and appropriate complexity gates. Statistical analysis was carried out using linear regression with a significance threshold of p ≤ 0.05 (GraphPad Prism, v. 10.6, Boston, MA, USA). Phenotypic structure and cell size determination assays were carried out using cultures at 30–40% confluence in T75 flasks (Corning, NY, USA), labeled with CellTracker™ Fluorescent Probes (Thermo-Scientific, Waltham, MA, USA) to mark both the cytoplasm and the plasma membrane, and DAPI (ThermoFischer, Waltham, MA, USAB) for nuclei labeling. Labeling analysis was carried out using CellProfiler^®^ software (Broad Institute, Cambridge, MA, USA), applying a cell mask pipeline for quantitative delimitation of cell size. Relative diameter analyses were carried out using GraphPad Prism (v. 10.6, Boston, MA, USA) and one-way ANOVA for statistical evaluation.

### 2.5. Semi-solid colony growth assay

For the semi-solid colony formation assay, the Hs578T_wt, Hs578T_ctr, and Hs578T_ko cell lines were subcultured into 6-well plates (Corning, NY, USA) at 1×10² cells per well (technical triplicates and biological duplicates). Prior to seeding, the 6-well plates were coating with UltraPure™ Agarose, High Melting (> 95°C) (Invitrogen, Thermo-Scientific, Waltham, MA, USA), sterilized, and used at a final concentration of 0.6% (1:1 agarose/DMEM with 20% FBS) to form the bottom “hard agarose” layer. Cells were then plated on top of the hard agarose layer, allowed to settle for 10 min before addition of the top layer of 0.35% agarose (1:1 agarose/DMEM with 20% FBS), referred to as “soft agarose”. After 21 days (with medium changes every four days), brightfield images were acquired (Invitrogen™ EVOS™ XL Core Imaging System, Fisher Scientific, Waltham, MA, USA), four images were acquired per symmetric field for each well (10× magnification). Colony number, diameter, and relative area were quantified using Python (*pandas*, *opencv*, and *math* packages), and statistical analysis was carried out using two-way ANOVA with a significance threshold of p ≤ 0.05 (GraphPad Prism, v. 10.6, Boston, MA, USA).

### 2.6. Wound Healing assay

For the wound healing (scratch) assay, the Hs578T_wt, Hs578T_ctr, and Hs578T_ko cell lines were plated onto 6-well plates (Corning, NY, USA) at 1×10⁴ cells per well (technical triplicates and biological duplicates) and grown to 80% confluence. All wells were then treated with Mitomycin (10 ng/mL, Sigma-Aldrich, St. Louis, MO, USA) for 24 h to inhibit cell proliferation. After the incubation period, a straight scratch was made at the center of each well using a 200 µL pipette tip, generating a cell-free gap. Cell migration toward the center of the scratch was monitored every 6 h over 24 h.

After 24 h, brightfield images were acquired (Invitrogen™ EVOS™ XL Core Imaging System, Fisher Scientific, Waltham, MA, USA), four images per symmetric field for each well (10× magnification). Migration analysis was carried out using ImageJ software (NIH, Bethesda, MD, USA) and the Wound Healing Size Tool plugin, and statistical analysis of relative migration area was conducted using two-way ANOVA with a significance threshold of p ≤ 0.05 (GraphPad Prism, v. 10.6, Boston, MA, USA).

### 2.7. Transwell invasion assay

The migratory potential of the Hs578T_wt, Hs578T_ctr, and Hs578T_ko cell lines was also assessed using Transwell™ inserts (8-µm pore size; BD Bioscience, San Jose, CA, USA). A total of 5×10⁴ cells were seeded onto the upper surface of the porous membrane in 300 µL of serum-free medium, using DMEM 0% FBS. The lower chamber received 700 µL of medium containing the corresponding chemoattractant (DMEM 10% FBS). Cells were incubated for 16–18 h at 37°C in 5% CO₂). After incubation, non-migrating cells were removed from the upper surface with cotton swabs. Inserts were fixed in cold methanol for 20 min and stained with Mayer’s hematoxylin for 20 min. After drying, migrating cells, which adhered to the lower surface of the membrane, were counted at 200× magnification using an EVOS FL Fluorescence Cell Imaging System (Fischer Scientific, Waltham, MA, USA). Four representative fields per insert were analyzed, with duplicate inserts for each condition.

### 2.8. Time-lapses and Cell track

The Hs578T_wt, Hs578T_ctr, and Hs578T_ko cell lines were subcultured into T75 culture flasks (Corning, NY, USA) at an approximate concentration of 1×10⁵ cells/mL. After subculturing and initial cell attachment, one field of view per culture flask corresponding to 20–30% confluence was selected. Using the CytoSMART™ System Live Cell Imaging platform (Lonza, Basel, Switzerland), images were acquired during 72 h, with images being captured every 5 min.

Image processing, including color inversion, adjustment of cell position, background correction, and removal of outlier frames (i.e., frames affected by lag in the Lonza imaging system), as well as analysis of cell displacement and the dynamics of cells and their associated vesicles, was carried out using ImageJ with the *TrackMate* plugin (with optimized detection settings: *DoG* detector, automatic threshold, and *LAP tracker*). After processing the final image sets, the video editing software KdenLive (v. 23.08.3, KDE, open-source, Berlin, Germany) was used to generate time-lapse videos (0.01 s per frame).

Analyses of cell displacement, dynamics, and associated statistics were performed using Python (*numpy*, *skimage*, and *tqdm* packages) in the Google Collab environment, with the following parameters applied to avoid overestimated cell movement values: *MIN_AREA = 10; MAX_DISPLACEMENT = 50; L_LENGTH = 30; and PROCESS_EVERY_N_FRAMES = 5*.

### 2.9. Migration qRT-PCR panel expression in Hs578T cell lines

The Hs578T_wt, Hs578T_ctr, and Hs578T_ko cell lines were cultured in medium-sized flasks (T75, Corning, NY, USA) until reaching 90% confluence. Total RNA was then extracted from 1×10⁶ cells using the Illustra RNAspin Mini Kit (Cytiva Life Sciences, Marlborough, MA, USA). cDNA synthesis was carried out with 2 µg of RNA from each experimental condition using the FireScript RT cDNA Synthesis Kit (Solis BioDyne, Tartu, Estonia). The yield and purity of the synthesized cDNA were assessed by the 280/260 nm ratio (280/260 ≥ 2.0), using a Nanodrop 1000 (Thermo-Scientific, Waltham, MA, USA), according to the manufacturer’s instructions.

For qRT-PCR, the Power SYBR™ Green Master Mix kit (Cat #4385612, ThermoFischer, Waltham, MA, USA) was used with a 40-cycle amplification protocol following the basic recommendations of the equipment (ViiA7 Real-Time PCR System, ThermoFischer, Waltham, MA, USA). Each reaction contained 2.5 µL of total cDNA, 2.5 µL of primers (forward + reverse, 400 nM) for the migration target genes (or 2.5 µL of nuclease-free H₂O in negative-control wells), and 5.0 µL of Power SYBR™ Green Master Mix (optimized volumes). The primers listed in the Table below (Table 2) were obtained from Exxtend – *Soluções em Oligos* (Paulínia, São Paulo, Brazil), lyophilized and desalted before use.

**Table 2:**
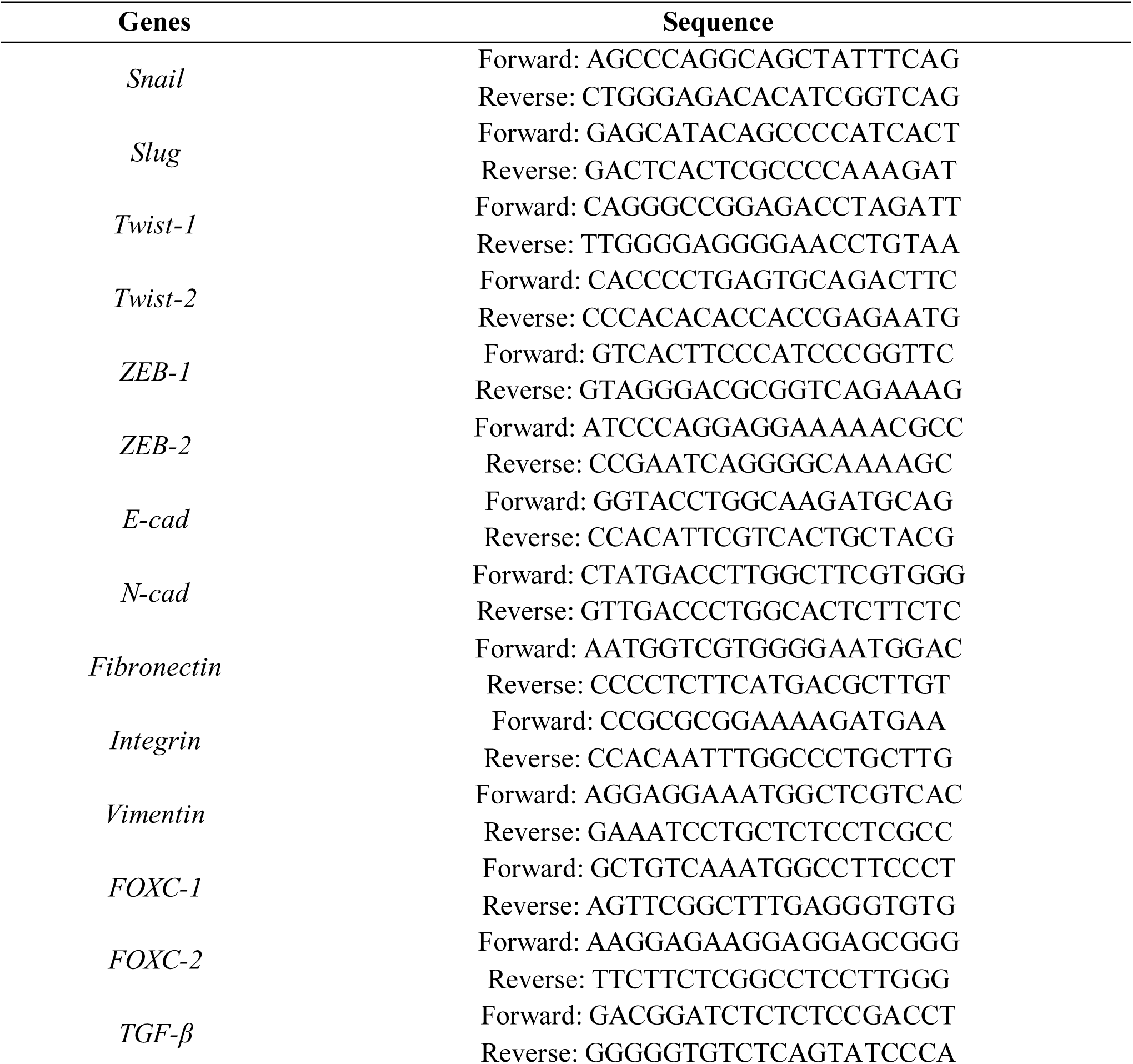
Primers sequences of the migration genes analyzed.

The references control genes used were: HSP90A1B (400 nM, Forward: CTCTGTCAGAGTATGTTTCTCGC; Reverse: GTTTCCGCACTCGCTCCACAAA), HPRT1 (400 nM, Forward: TGAGGATTTGGAAAAGGGTCT; Reverse: GAGCACACAGAGG GCTACAA), and HMBS (400 nM, Forward: TGGACCTGGTTGTTCACTCCTT; Reverse: CAACAGCATCATGAGGGTTTT).

All qRT-PCR assays were carried out in technical triplicates and biological duplicates. The mean Ct values of the internal control genes were used to calculate log₂FC values for each gene analyzed using the 2^−ΔΔCt^ method (relative to the reference cell line: MCF10A). The final results were statistically analyzed using two-way ANOVA with a significance threshold of p ≤ 0.05 (GraphPad Prism, v. 10.6, Boston, MA, USA).

## 3. Results

Identification of the lncRNA LINC01133 as a molecule of interest in TNBC was based on analysis of gene expression at the RNA level in a panel of seven non-tumorigenic and tumorigenic mammary cell lines. The MCF10A cell line was used as a non-tumorigenic reference cell line, to evaluate the upregulated or downregulated expression of several lncRNAs of known relevance in different molecular subtypes of breast tumor cell lines.

The results, shown in Figure 1, demonstrate upregulated expression of only a few lncRNAs, namely: BANCR, GHET1, and TERC, in the SK-BR-3, ZR-75-1, and MDA-MB-231 cell lines, respectively. However, LINC01133 stood out with the highest expression level, displaying a log₂FC close to 5, corresponding to approximately a 30-fold increase in expression in Hs578T cells, when compared with normal MCF10A and MCF12A and other tumor cell lines.

**Figure 1:**
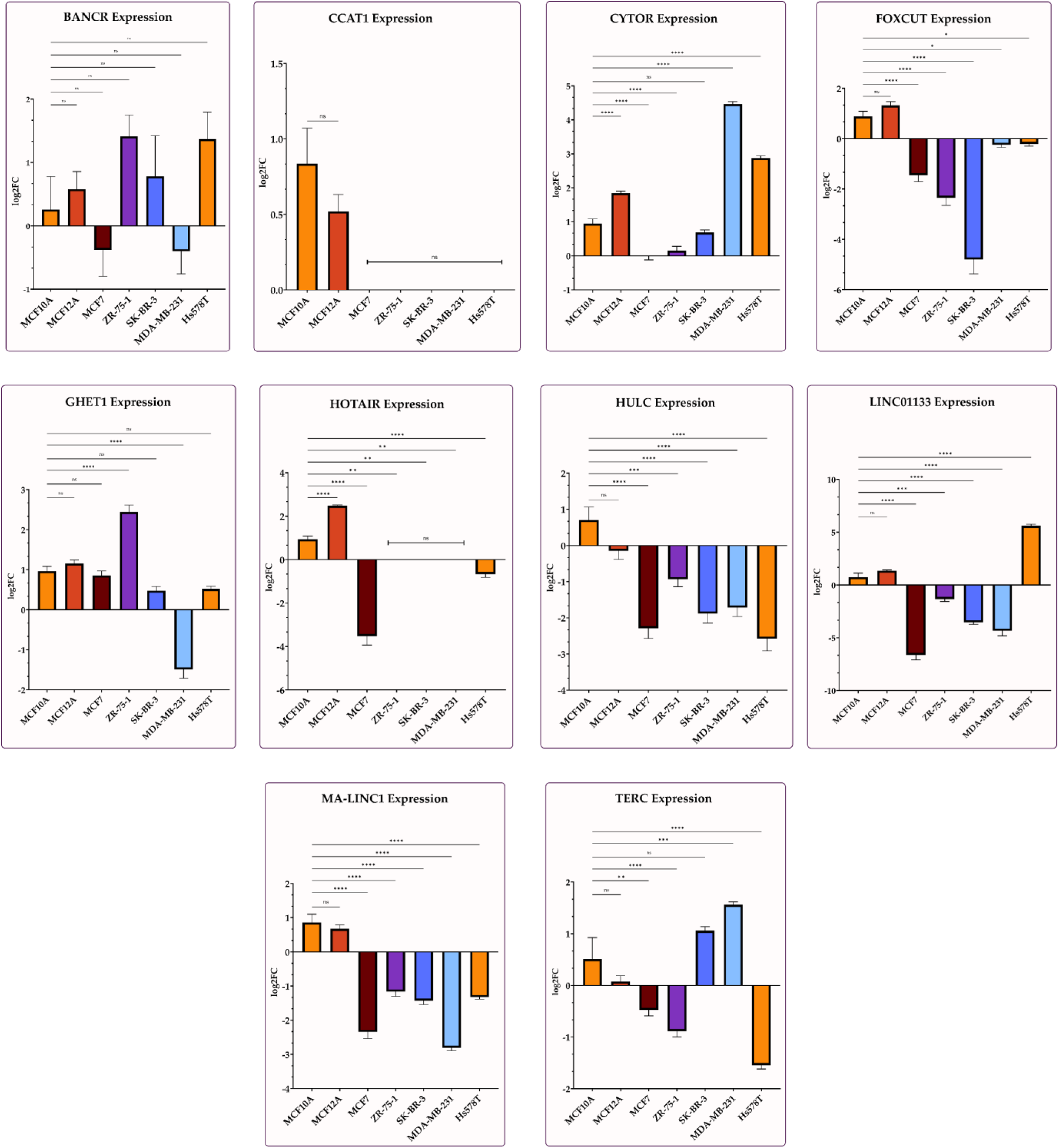
Transcriptional expression levels of lncRNAs of interest in a panel of non-tumorigenic (MCF10A and MCF12A) and tumorigenic mammary cell lines. Expression values are shown as log₂FC. Statistical analysis was carried out using two-way ANOVA (p-value: * ≤ 0.05; ** ≤ 0.01; *** ≤ 0.001; **** ≤ 0.0001; ns: non-significant). Analyses were based on technical and biological triplicates using GraphPad Prism v. 10.6.

Based on this observation, a genomic characterization of the LINC01133 gene was pursued aiming at genetic manipulation of the locus containing its exons. Although more than 30 genomic isoforms are annotated, the consensus structure comprises three canonical exons, which were considered for the design of gRNAs to enable CRISPR/Cas9-mediated knockout [9,15]. Lentiviral plasmids co-expressing the Cas9 enzyme and the gRNAs in cis were used to achieve complete knockout of the exonic regions. The first 250 nucleotides (5′ end) and the last 250 nucleotides (3′ end) of the LINC01133 gene were selected, with the primer sequences shown in Figure 2. The knockout procedure, involving lentiviral vector production in 293T cells, followed by transduction of the Hs578T_wt cell line and puromycin selection, was successfully completed, generating stable knockout cell lines as well as the Hs578T_ctr control line, which underwent the same protocol.

**Figure 2:**
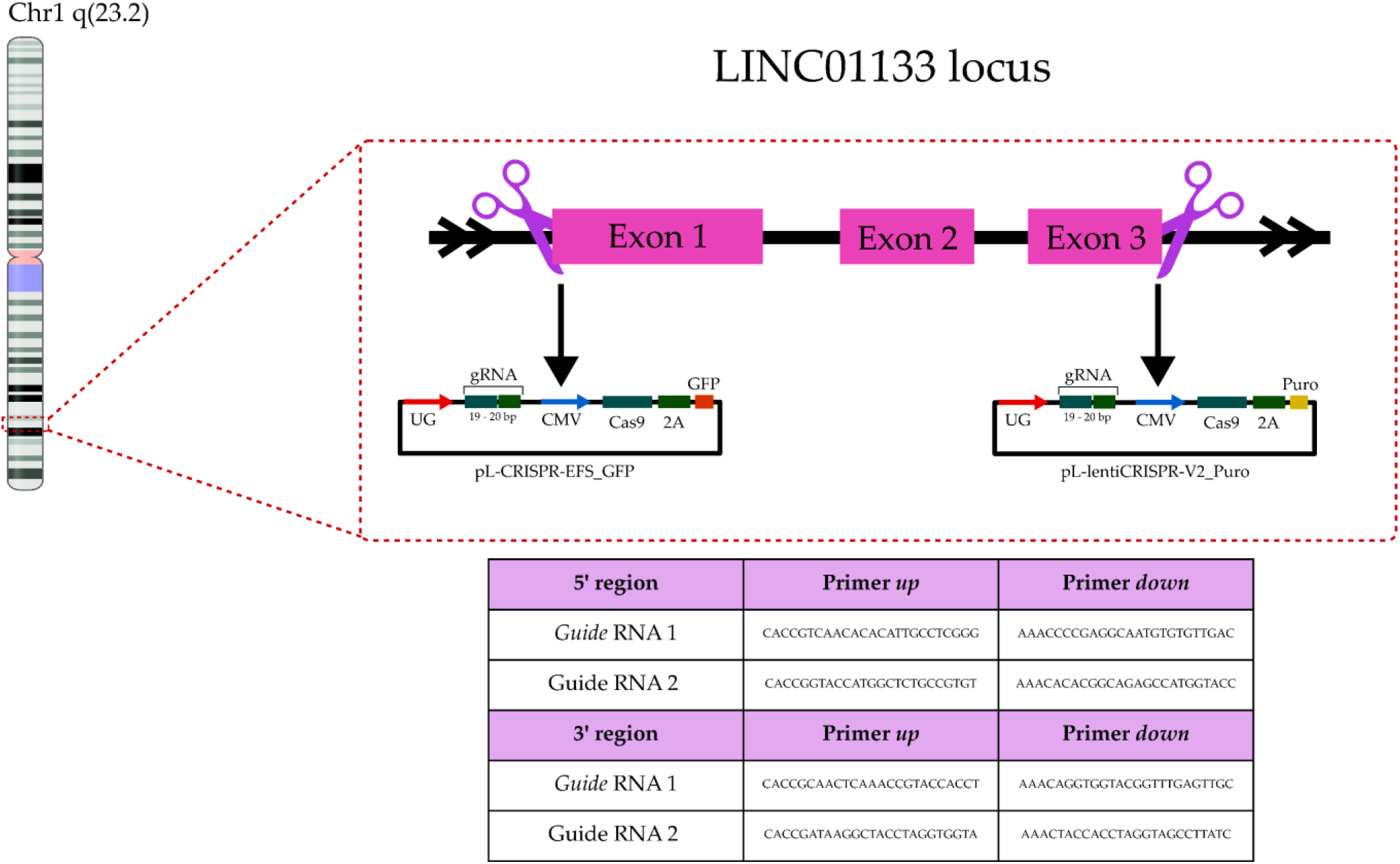
Schematic representation of the LINC01133 gene knockout process carried out in the Hs578T_wt cell line, highlighting the cut regions and the gRNA sequences used.

Upon establishment of this Hs578T cell line panel, phenotypic and molecular assays were undertaken. Morphological analyses using CellTracker™ fluorescence revealed an interesting effect of the LINC01133 gene knockout, namely a reduction of both cell diameter and area, as shown in Figure 3A. Growth curves under serum deprivation conditions indicated a limited impact of the knockout on cell proliferation parameters (Figure 3B), although in quantitative analyses the Hs578T_ko cell line showed a tendency for a shorter doubling time under moderate serum deprivation, as shown in Table 3. A substantial change in cell proliferation rate was observed for the genetically modified lines under serum deprivation (1% FBS), as shown in Figure 3B.

**Figure 3:**
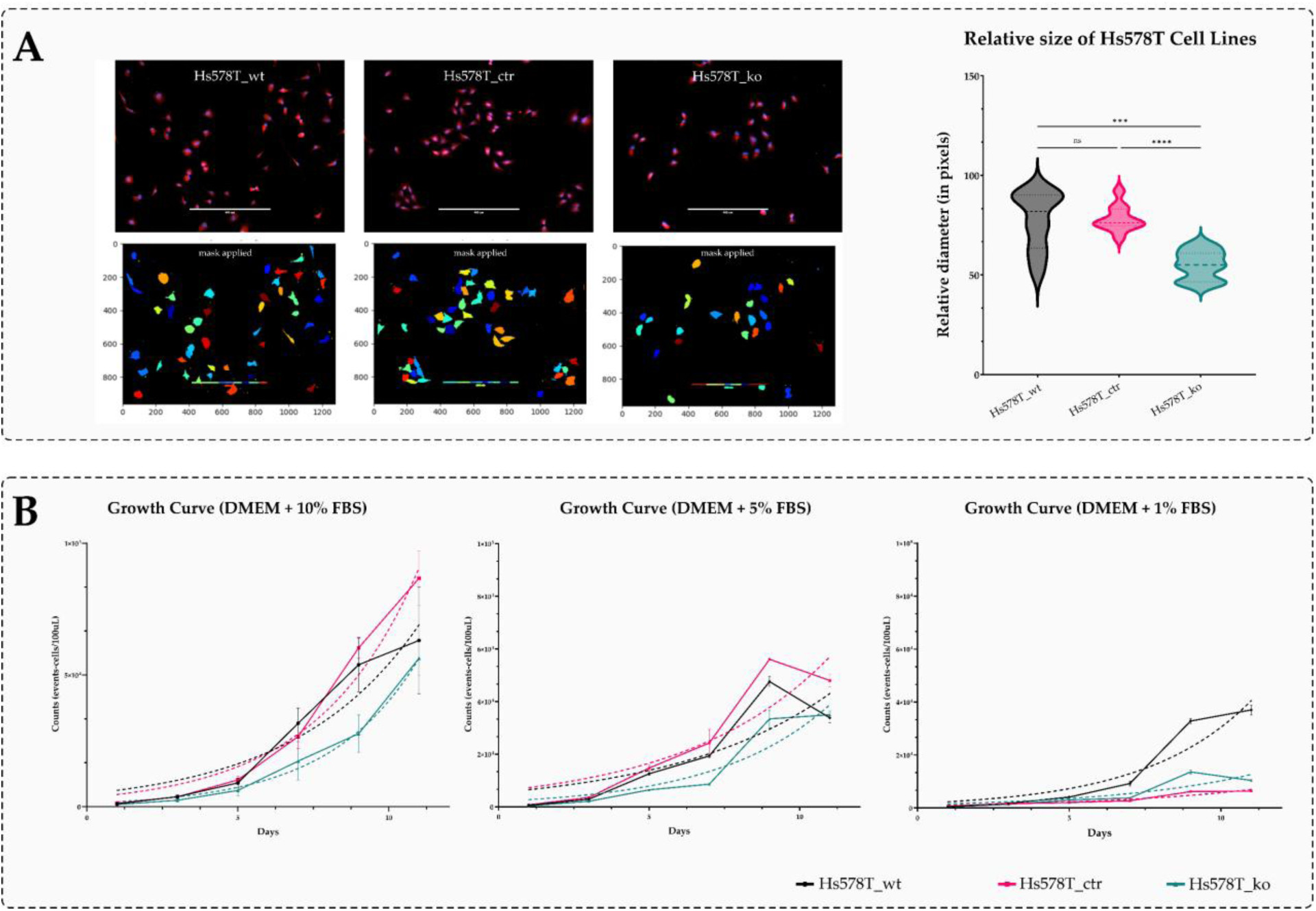
**A.** Morphological analysis of phenotypic structure of Hs578T_wt, Hs578T_ctr, and Hs578T_ko cell lines, using CellTracker™ Probes (red) and DAPI for nuclei labeling (blue); the diameter of cells was analyzed using CellProfiler for cell masking and one-way ANOVA for statistical analysis (GraphPad v. 10.6; p-value: *** ≤ 0.001; **** ≤ 0.0001; ns: non-significant). **B.** Growth curves of the three analyzedcell lines, undernormal andserum-deprivationconditions (DMEM + 10% FBS; DMEM + 5% FBS; DMEM + 1% FBS). Analyses were performed by flow cytometry (Accuri C6 Flow Cytometer, BD Biosciences, San Jose, CA, USA), and results were analyzed using GraphPad Prism v. 10.6 (linear regression with SEM).

**Table 3.** Comparative doubling time among Hs578T cell lines.

| Doubling Time | Hs578T_wt | Hs578T_ctr | Hs578T_ko | Difference Hs578T_ko (%) |
| --- | --- | --- | --- | --- |
| **DMEM + 10% FBS** | 1.875 to 4.619 | 1.913 to 2.844 | 1.473 to 2.813 | 34.0% shorter |
| **DMEM + 5% FBS** | 2.218 to 7.020 | 2.224 to 5.480 | 1.791 to 3.912 | 38.3% shorter |
| **DMEM + 1% FBS** | 1.951 to 3.035 | 3.055 to 4.443 | 2.402 to 4.774 | 43.9% higher |

In contrast to the cell proliferation data, cell migration and invasion assays revealed striking effects of lncRNA01133 knockout. In agarose semi-solid medium not only a higher number of colonies were formed by Hs578T_ko cells after 21 days (Figure 4), but these colonies also exhibited larger absolute dimensions, in terms of both area and diameter.

**Figure 4:**
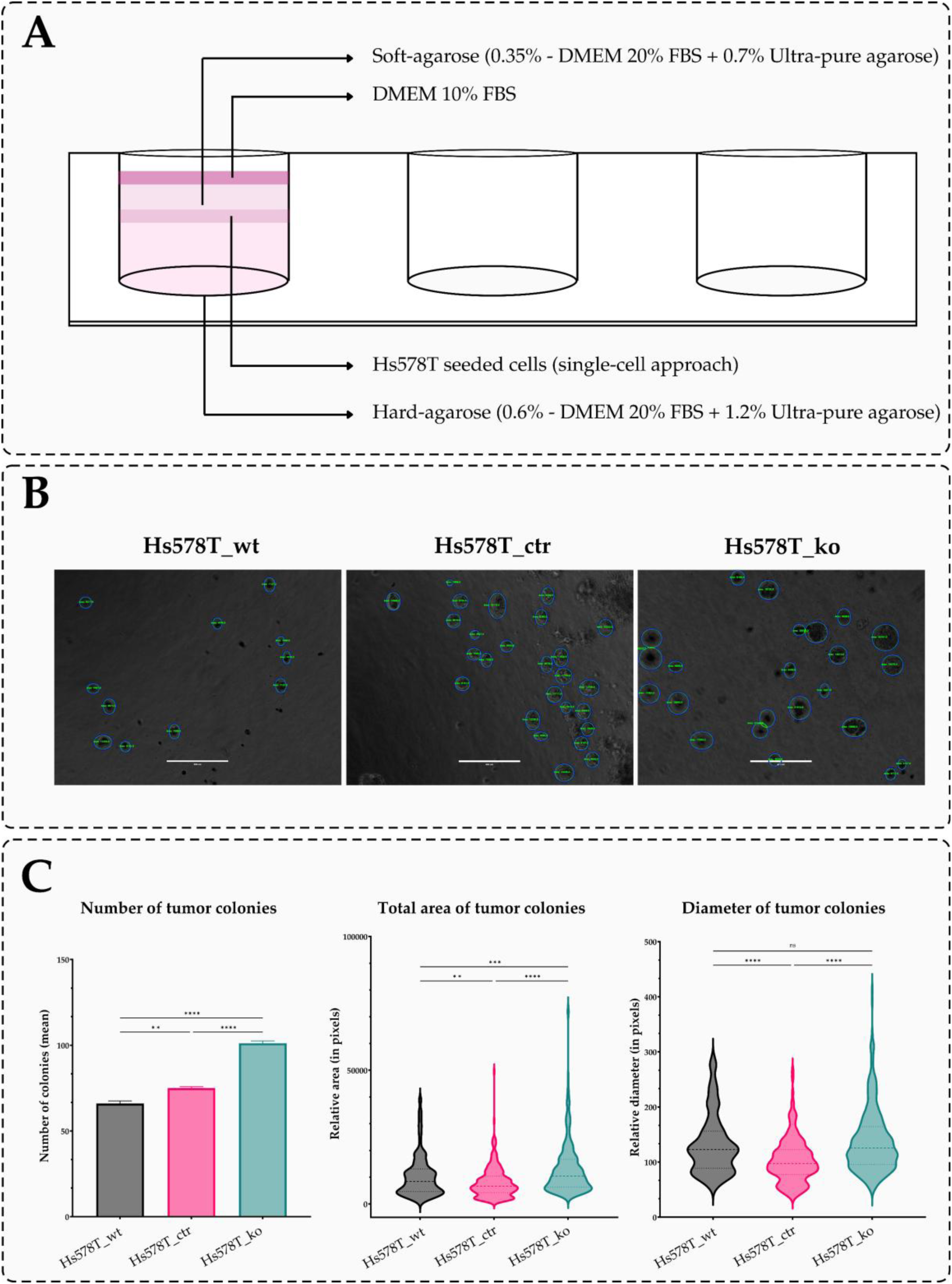
Growth in semi-solid agarose medium. A. Schematic representation of cell plating between agarose layers; **B.** Representative images of the identified colonies using *pandas*, *opencv*, and *math* packages (Python IDE); **C.** Results of absolute colony counts and their respective diameters and areas, analyzed using GraphPad Prism v. 10.6 (two-way ANOVA, p-value: ** ≤ 0.01; *** ≤ 0.001; **** ≤ 0.0001; ns: non-significant).

In wound healing experiments, which allowed visualization of bidimensional cell migration following a scratch introduced at the center of the well, a significantly increased migratory capacity was also observed in the Hs578T_ko line. Specifically, the knockout cells showed an approximately 10-fold increase in relative wound closure of the scratched area compared with the Hs578T_wt and Hs578T_ctr lines. This effect is particularly notable given the use of mitomycin, a potent antiproliferative agent that prevents false-positive results arising from cell proliferation within the scratched region [16] (Figure 5).

**Figure 5:**
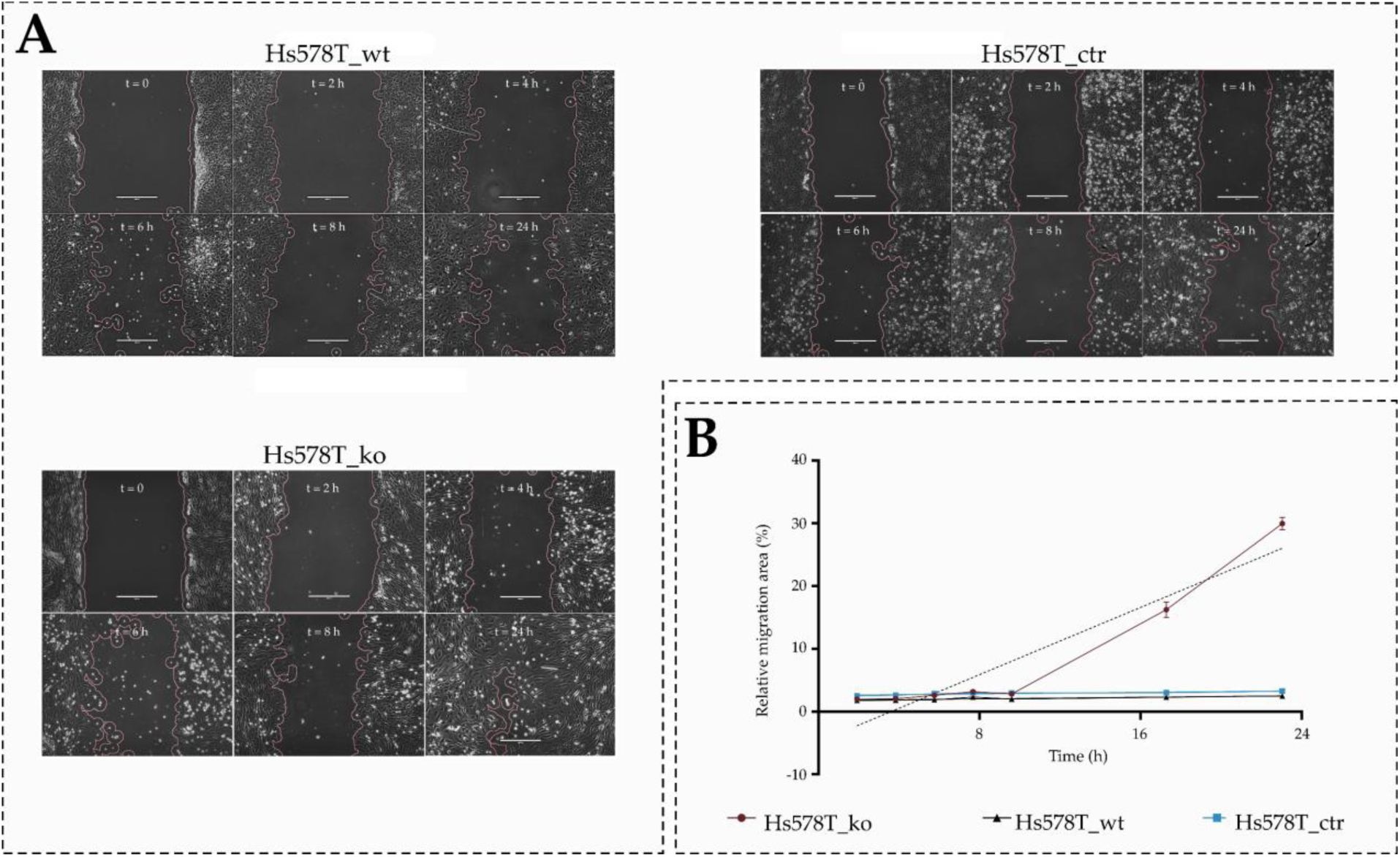
Wound healing assay of Hs578T_wt, Hs578T_ctr, and Hs578T_ko cell lines. **A.** Representative images of the scratch and cell migration over the 24h assay period; **B.** Comparative statistical analysis (plot) of relative migration of all three Hs578T cell lines, analyzed using GraphPad Prism v. 10.6 (linear regression with tendency line with SEM).

The enhanced migratory phenotype of the Hs578T_ko line was further supported using the Transwell assay (Figure 6). In this system, the knockout cell line displayed a markedly increased migratory capacity, with the migration level through the porous membrane being approximately 55-fold higher than that observed for the Hs578T_wt and Hs578T_ctr cell lines.

**Figure 6:**
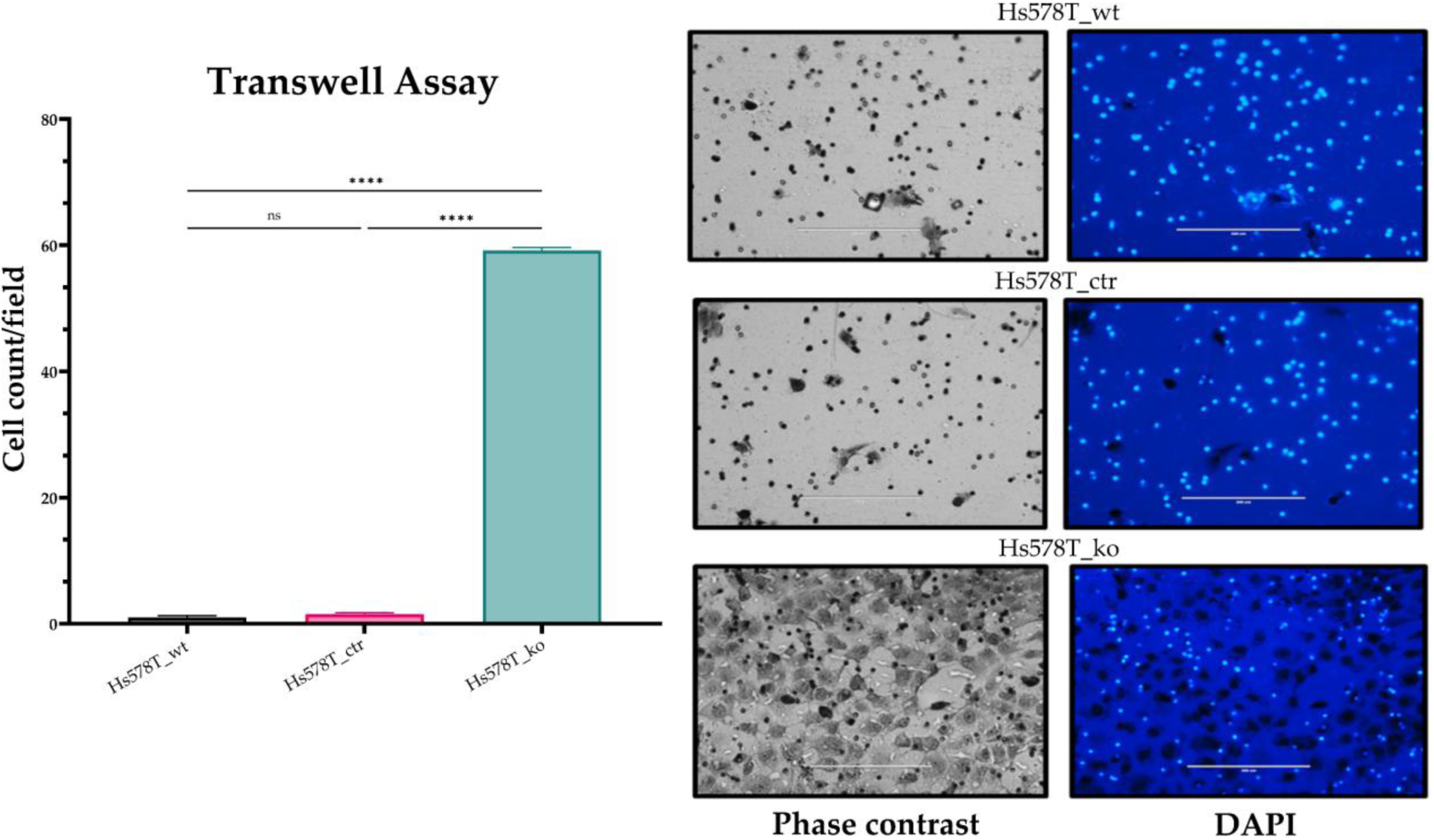
Transwell migration and invasion assays of the Hs578T_wt, Hs578T_ctr, and Hs578T_ko cell lines analyzed. The results were analyzed by one-way ANOVA followed by Bonferroni post-hoc comparison (p-value: **** ≤ 0.0001; ns: non-significant).

To enable a comparative analysis of cell dynamics across the established cell panel, experiments were designed to investigate the migratory and invasive properties associated with LINC01133 knockout. Static-field time-lapse imaging combined with cell-tracking analysis was performed in Hs578T_wt, Hs578T_ctr, and Hs578T_ko cell lines.

Time-lapse microscopy revealed distinct differences in overall cell displacement dynamics among the three Hs578T-derived cell lines. Gaussian displacement distribution analysis showed a higher proportion of cells with elevated migratory potential in the Hs578T_ko cell line. Among the 50 cells analyzed, the knockout line presented the highest number of cells with effective displacement above 600 pixels, as well as increased off-axis movement. In addition, time-lapse imaging revealed an increased movement of vesicular particles in the knockout cells.

As shown in Figure 7A and in the videos provided in Supplementary Material S1, the Hs578T_ko cell line exhibited greater motility and dynamic behavior, while the parental and control lines displayed more pronounced rotational axis dynamics. Quantitative analysis of cell movement (Figures 7B and 7C), with statistical outcomes summarized in Figure 7D, demonstrated an average displacement approximately 15% higher in the Hs578T_ko cell line, with differences becoming more evident after two days of assay progression. Furthermore, the displacement distribution shifted toward higher values, accompanied by a more homogeneous cellular density in this region.

**Figure 7:**
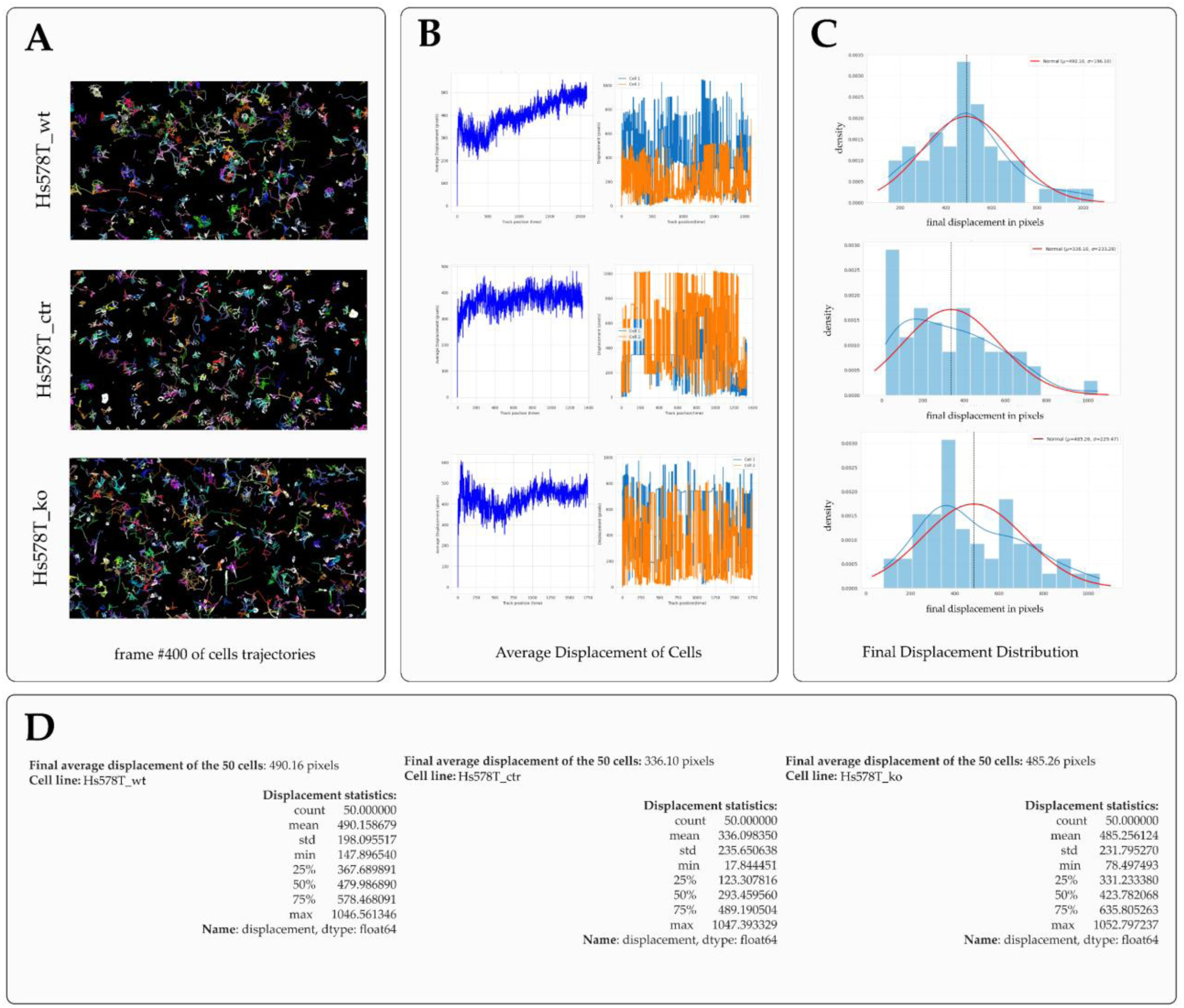
Cell-tracking assay carried out with the Hs578T_wt, Hs578T_ctr, and Hs578T_ko cell lines. **A.** Visual representation of the trajectory calculations analyzed over 72h; **B.** Cellular average displacement and representative individual displacement over time (pixels); **C.** Final displacement distribution by pixels versus cellular density; **D.** Quantitative comparative statistical results. A total of 50 cells per brightfield were analyzed.

Based on the phenotypic evidence described above, a transcriptional expression panel of genes associated with migratory processes was assembled to initiate a molecular investigation of the mechanisms predominantly affected by the LINC01133 knockout. As shown in Figure 8, significant differences in log₂FC expression were observed for several classical genes involved in: ECM establishment and degradation, cell adhesion, EMT, and cell migration. Among the 14 genes analyzed, those encoding integrins, fibronectin, and vimentin were particularly notable, showing upregulated expression in the Hs578T_ko line. In contrast, genes such as TWIST2 and ZEB1 were also significantly downregulated in the LINC0113 knockout.

**Figure 8:**
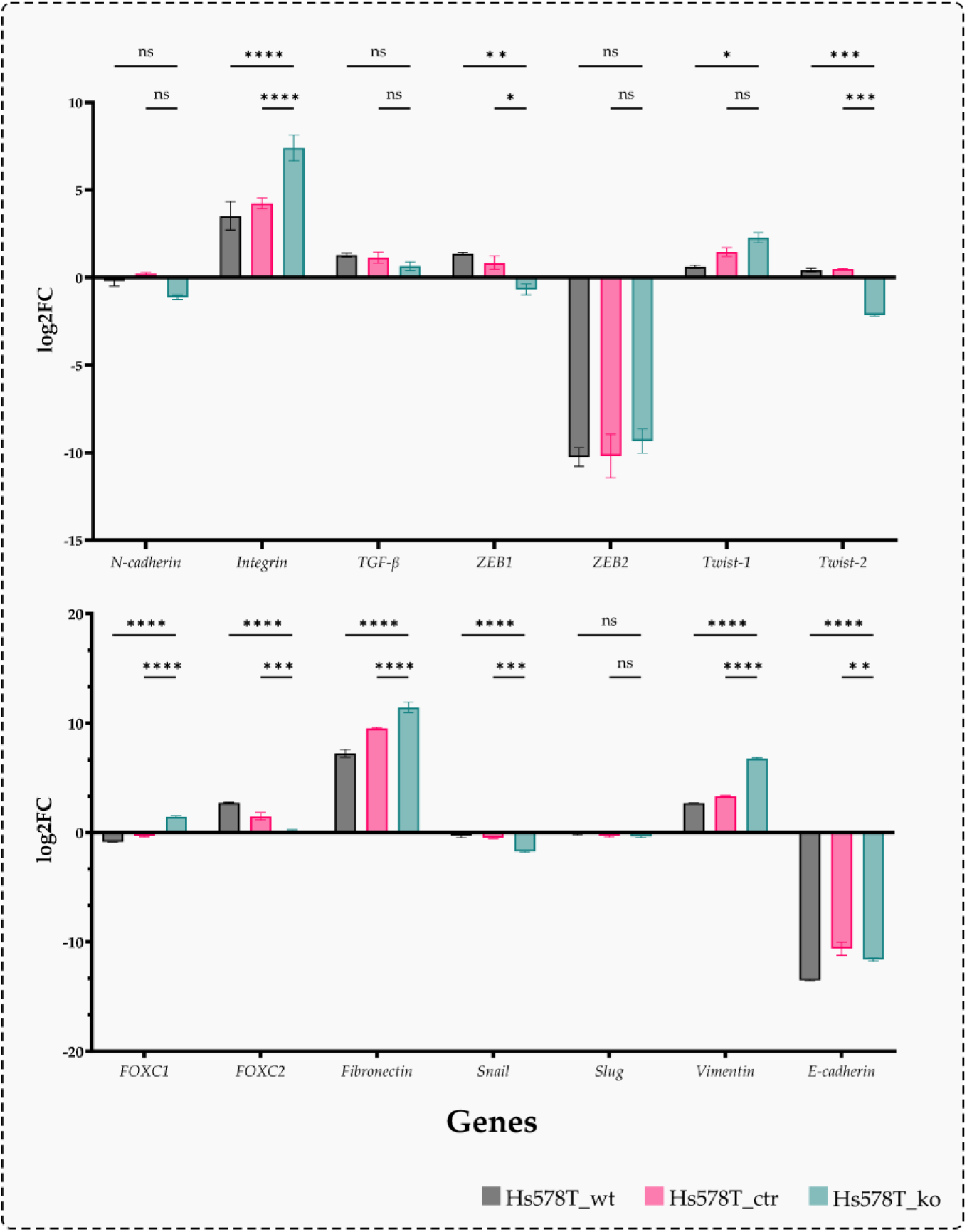
Comparative transcriptional expression panel of genes associated with cell migration processes in the three analyzed Hs578T cell lines. Expression values are shown as log₂FC. Statistical analysis was performed using two -way ANOVA (p-value: * ≤ 0.05; ** ≤ 0.01; *** ≤ 0.001; **** ≤ 0.0001; ns: non-significant). Analyses were based on technical quadruplicates and biological duplicates, using GraphPad Prism v. 10.6.

## 4. Discussion

Over the last 20 years, lncRNAs have become established as molecules of interest across multiple tumor types, performing diverse functions that may be tumor-suppressive or oncogenic, within the tumor microenvironment and in the intratumoral environment [17]. Long known transcripts such as HOTAIR, MALAT1, TUG1, FOXCUT, among others, have been widely studied as potential predictive markers and/or therapeutic targets in Oncology, which further increases interest in lncRNA research in Clinical Medicine [18–20]. Although LINC01133 is still relatively understudied, it has been investigated as a potential tumor suppressor since 2017, with evidence linking its function to pluripotency proteins, such as KLF4 and SOX4, as well as to possible roles in cancer stem cell–related mechanisms and modulation of metastatic processes [10,11,21].

The complexity of the dual relationship between lncRNAs and cancer is well illustrated in breast tumors. Despite the classical molecular classification into five tumor subtypes (Normal-like, Luminal A, Luminal B, HER2+ – including the HER2-enriched subclass – and Basal-like), breast cancer heterogeneity is extensive enough to define more than 10 molecular subtypes and/or subclasses [22,23]. Each of these is characterized by a complex molecular network that may be associated with uncontrolled proliferation, enhanced migration, increased metastatic potential, or, in more specific contexts, heightened inflammatory capacity and immune evasion [24].

When specifically focusing on triple-negative subtype, this high level of complexity becomes particularly evident. These features, mainly the asymptomatic progression and high capability of migration, invasion and distant metastasis, clearly hinder the identification of molecules with broad clinical applicability [25]; however, the discovery of clinically relevant molecules in well-established cell lines can open new avenues for translational research, enabling their potential application, as suggested here for the lncRNA LINC01133.

In the results presented in Figure 1, the upregulated differential expression of LINC01133 was restricted to the Hs578T cell line, in contrast to the downregulated expression observed in the MDA-MB-231 cell line, both cell lines being representatives of the TNBC class. This difference likely reflects the heterogeneity among TNBC subclasses across these cell lines. Nevertheless, it is evident that, depending on the expression of other key molecules, LINC01133 is associated with modulatory processes of tumor progression. Importantly, its significantly increased expression in the Hs578T cell line was a key factor guiding the selection of this model for LINC01133 gene knockout experiments.

The LINC01133 knockout strategy was designed to avoid off-target effects that could generate false-positive phenotypic or molecular results. gRNAs were selected to ensure effective knockout, particularly considering the nature of the target gene—an intergenic lncRNA whose biological function may be associated with the mature transcript, the genomic locus itself, or the act of transcription. In this context, the experimental strategy prioritized targeting critical regions of the locus to robustly abolish LINC01133 transcription without interfering with regulatory elements of neighboring genes [26] (Figure 2). In addition, knockout validation was performed using independent transcriptional analyses, ensuring consistent reduction of lncRNA expression and minimizing interpretations driven by nonspecific effects. Generation of a genetically modified control cell line (Hs578T_ctr), carrying an inserted empty plasmid backbone, was also essential to provide an additional control for subsequent assays, thereby excluding false positives resulting from genetic manipulation rather than from the knockout itself.

Following the knockout process, subsequent experiments focused on comparative analyses of phenotypic and molecular characteristics among the knockout, parental, and control cell lines. The first assay evaluated the single-cell structural features of the Hs578T_wt, Hs578T_ctr, and Hs578T_ko lines, given the relevance of relative tumor cell diameter in proximal and distal metastasis. Smaller tumor cells often exhibit higher migratory and invasive potential, which is associated with poorer oncological prognosis [27,28]. CellTracker™ fluorescence assays revealed that the average cell diameter of the Hs578T_ko line was approximately 31% smaller than that of the Hs578T_wt and Hs578T_ctr lines (Figure 3A). Together with the phenotypic and molecular findings presented in the Results section, these data indicate that the knockout affects not only migration-related mechanisms, but, also, cell morphology, suggesting a modulatory role in invasive tumor processes.

Regarding growth curve assays, under adequate serum supply conditions (DMEM 10% FBS and DMEM 5% FBS), all cell lines exhibited similar growth patterns. In contrast, under low serum conditions (DMEM 1% FBS), only the Hs578T_wt line maintained a high proliferative rate (compared to the other cell lines) (Figure 3B). These results suggest that the genetic modifications introduced in Hs578T_ctr and Hs578T_ko strongly affect adaptative responses to energetic stress, impairing proper metabolic adaptation. Since this effect was observed in both modified lines, it cannot be directly attributed to the LINC01133 knockout. Therefore, the growth curve data indicate that LINC01133 knockout does not significantly alter the tumor proliferative activity, the observed increase in malignancy being more closely linked to other mechanisms of tumor progression.

Semi-solid agarose growth assays, together with comparative morphology data, further highlight the impact of LINC01133 knockout on migration and invasion mechanisms. The single-cell seeding approach was important to demonstrate both a higher absolute number of colonies and increase of relative colony area and diameter in the Hs578T_ko line. These findings suggest that LINC01133 knockout promotes more aggressive phenotypic patterns in this TNBC model, characterized by smaller cells that nonetheless organize into more numerous and larger colonies (Figure 4). In the context of LINC01133 as a tumor suppressor, these results illustrate how its role may be multifaceted. Although previous studies, together with the present data, support a tumor-suppressive potential of LINC01133 in TNBC, it is important to consider that depending on its expression levels, LINC01133 may be more associated to a fine tuning rather than to a pure dual (oncogenic or tumor suppressor) effect (10,11,13,22,30). This effect may not occur when expression is reduced to negligible levels, as under knockout conditions. Consequently, LINC01133 loss clearly increases malignancy, although, from an evolutionary perspective, excessive malignancy may reduce tumor fitness and significantly shorten patient survival. The evolutionary fitness framework in Oncology remains underexplored, but is essential for under-standing tumor progression and prognostic complexity.

Consistent with these observations, wound healing and Transwell assays further consolidate the link between LINC01133 knockout and increased malignancy through enhanced migratory and invasive mechanisms. Scratch assays clearly demonstrated a strong association between LINC01133 and cell migration, likely mediated by modulation of intracellular components such as membrane and cytoskeletal proteins, as well as factors related to the intra-tumoral environment that stabilize tumor mass architecture and interact with ECM components [29–31]. The introduction of Mitomycin treatment in the wound healing assay was critical to separate cell migration from cell proliferation (Figure 5). Transwell assays further showed how increased migratory activity, combined with smaller cell size, confers higher invasive potential (Figure 6). Breast tumors typically metastasize first to axillary lymph nodes before disseminating to distant organs, such as lungs, liver, bones, and brain [32,33]. Smaller cells with enhanced migratory capacity may initiate these processes more rapidly and efficiently in vivo, as suggested by this in vitro model.

Based on cell tracking assays, the increased displacement, off-axis movement, and higher proportion of highly migratory cells observed in the Hs578T_ko cell line indicate that LINC01133 knockout is associated with enhanced migratory and invasive behavior. The shift toward a Brownian-like migration pattern suggests not only increased motility but also a greater capacity for dynamic cell – cell interactions, which may contribute to a more aggressive cellular phenotype [34].

Although their molecular content remains undefined, the elevated presence and movement of vesicular particles in the knockout line, may be functionally linked to the enhanced migratory and invasive properties observed [35]. Together, these findings support a possible role for LINC01133 in restraining cell motility, its loss appearing to promote dynamic behaviors associated with tumor progression.

Considering the strong phenotypic evidence linking LINC01133 knockout to migration and invasion, a transcriptional expression panel of genes associated with these processes was assembled for the three Hs578T cell lines analyzed, using MCF10A as a normal/non-tumorigenic control. qRT-PCR analyses revealed transcriptional changes consistent with a more aggressive phenotype in the Hs578T_ko line, displaying expression profiles clearly distinct from those of the Hs578T_wt and Hs578T_ctr lines.

Among the analyzed genes, selective induction of FOXC1 was observed in the Hs578T_ko line, in contrast to its repression in the Hs578T_wt and Hs578T_ctr lines. This finding is particularly relevant given FOXC1 involvement in tumor aggressiveness and cell migration. Concurrently, a reduction in FOXC2 expression was observed, suggesting coordinated transcriptional regulation within the FOX family and a functional reorganization of EMT and invasion-related circuits [36].

Analysis of structural and ECM-interaction markers reinforced this scenario. Fibronectin expression was significantly higher in Hs578T_ko cells, indicating increased ECM remodeling and altered adhesion dynamics. In parallel, increased vimentin expression supported a shift toward a mesenchymal phenotype. Transcriptional overexpression of integrins further supported these findings, pointing to enhanced ECM anchorage efficiency and exacerbated migratory and invasive phenotypes [37].

Classical EMT regulators displayed a partially noncanonical pattern consistent with a state of high cellular plasticity. Snail and Slug expression suggested selective activation of EMT axes, whereas the observed inhibition of N-cadherin in the knockout line indicates that increased malignancy may not strictly depend on the classical E-cadherin to N-cadherin switch. Instead, it may involve collective migration mechanisms driven by reorganization of intracellular proteins and cell–cell and cell–ECM interaction components [38].

Finally, analysis of ZEB and TWIST factors revealed a distinct regulatory balance in the knockout line. While ZEB1 and TWIST2 expression was reduced in Hs578T_ko cells, TWIST1 expression increased progressively and reached its highest level under knockout conditions. This pattern suggests that loss of LINC01133 favors invasive pathways predominantly mediated by TWIST1, independently of ZEB1 or TWIST2 activation. Together, qRT-PCR data support the notion that LINC01133 knockout induces directed transcriptional reprogramming that enhances migration and invasion through selective transcription factor activation, ECM remodeling, and strengthened cell–matrix interactions, culminating in a more malignant phenotype [39,40].

Overall, the phenotypic and molecular assays presented here demonstrate a strong influence of LINC01133 activity in TNBC. Its basal expression in the Hs578T_wt line is likely associated with fine modulation of tumor progression–related processes, a hallmark of TNBC in vivo, being characterized by high malignancy and clinical silence [41]. Unlike studies that frame LINC01133 strictly as a tumor suppressor or oncogenic agent, the present findings support a dual role in tumor contexts. Its overexpression may shift biological activity toward a dominant suppressive role, although further studies are required to confirm this hypothesis. Nevertheless, LINC01133 emerges as a highly relevant molecule capable of intricately modulating molecular pathways associated with migration and invasion, which are primary targets in oncological prediction and therapeutic strategies for breast cancer.

## 5. Conclusions

The results of this study consistently demonstrate that deletion of the lncRNA LINC01133 triggers a phenotypic and transcriptional reprogramming that favors migratory and invasive pathways in the Hs578T cell line. Loss of LINC01133 promotes a marked increase in cell motility, transmembrane invasion, and anchorage-independent colony formation, accompanied by morphological changes compatible with enhanced cellular plasticity, without inducing a proportional increase in basal proliferation.

At the molecular level, LINC01133 loss is related with the selective activation of genes involved in extracellular matrix remodeling, cell adhesion, and EMT-related regulatory axes, suggesting that this lncRNA functions as a fine modulator of pro-invasive transcriptional networks rather than as a binary regulator of classical EMT. This profile clearly shows the concept that TNBC progression may be driven by intermediate states of cellular plasticity that are highly migratory and adaptative.

Collectively, these findings identify LINC01133 as a potential regulator of migratory phenotype in TNBC and provide a mechanistic basis for its future exploration as a potential biomarker of invasive progression and/or as a target in combinatorial therapeutic strategies. Further studies, including in vivo validation and analyses in clinical cohorts, will be essential to establish the translational relevance of LINC01133 in the context of TNBC tumor heterogeneity and its metastatic evolution.

## Supporting information

Trajectories outputs of the Hs578T cell lines.

## Author Contributions

Conceptualization, LTJ, HCJF, ACOC, MCS.; Data curation, LTJ, HCJF.; Formal analysis, LTJ, HCJF.; Investigation, LTJ, HCJF.; Methodology, LTJ, HCJF, ACOC, MCS.; Visualization, LTJ.; Validation, LTJ, HCJF.; Writing – Original Draft, LTJ.; Writing – Review and Editing, LTJ, HCJF, ACOC, MCS.; Funding Acquisition, MCS.; Supervision, MCS, ACOC.; Project Administration, MCS.

## Institutional Review Board Statement

The use of genetically modified cell lines (Hs578T_ctr, and Hs578T_ko) was authorized by the Internal Biosafety Committee of the University of São Paulo Medical School (FM-USP) and the São Paulo Cancer Institute (ICESP) under process CTNBio 7476/2021, following approval for the generation of knockout cell lines using the lentiviral system at the NB2 facility.

## Informed Consent Statement

Not applicable.

## Data Availability Statement

The raw data used and/or analyzed during the current study are available in the BioStudies Database, under the following DOI: 10.6019/S-BSST3176. The dataset can be accessed at: https://www.ebi.ac.uk/biostudies/studies/S-BSST3176

